# The brain of the beholder: honouring individual representational idiosyncrasies

**DOI:** 10.1101/015594

**Authors:** Ian Charest, Nikolaus Kriegeskorte

## Abstract

In the early days of neuroimaging, brain function was investigated by averaging across voxels within a region, stimuli within a category and individuals within a group. These three forms of averaging discard important neuroscientific information. Recent studies have explored analyses that combine the evidence in better-motivated ways. Multivariate pattern analyses enable researchers to reveal representations in distributed population codes, honouring the unique information contributed by different voxels (or neurons). Condition-rich designs more richly sample the stimulus space and can treat each stimulus as a unique entity. Finally, each individual’s brain is unique and recent studies have found ways to model and analyse the interindividual representational variability. Here we review our field’s journey towards more sophisticated analyses that honour these important idiosyncrasies of brain representations. We describe an emerging framework for investigating individually unique pattern representations of particular stimuli in the brain. The framework models stimuli, responses and individuals multivariately and relates representations by means of representational dissimilarity matrices. Important components are computational models and multivariate descriptions of brain and behavioural responses. These recent developments promise a new paradigm for studying the individually unique brain at unprecedented levels of representational detail.

---

We would like to understand “the brain”. However, every brain is different. Our unique brains, products of our genes and individual experience, give rise to our unique personalities. Even the same person’s brain is constantly in flux, with its plasticity adapting the individual to a changing environment [1, 2, 3]. Given the idiosyncratic and plastic nature of any individual brain, it is amazing that brain science has been quite successful with a research paradigm that assumes that all brains are identical. At the gross scale of the global layout of functional regions, brain imaging studies have documented a significant degree of consistency across individual brains in both the functional decomposition and the approximate localisation of functional components. However, we know that functional correspondency between individual brains must break down at some level [4]. As far as we know, no neuron in a higher level cortical region in one person’s brain has an exact functional equivalent in another person’s brain.

In this paper, we argue that neuroscience needs to honour the uniqueness of the individual brain, the unique contribution of each patch of an individual cortex (and ultimately each neuron) to brain representations and processing and the particular properties and meaning of each particular stimulus (Figure 1). Most studies in neuroimaging have averaged across individuals, across cortical columns (within functional regions) and across particular stimuli. Reducing the complexity by averaging has provided a natural starting point for our investigations. In the domain of vision, early functional magnetic resonance imaging (fMRI) reported several cortical regions that selectively respond to particular categories of images. For example [5], contrasted brain activations elicited by faces and other objects and analysed the fMRI signal averaged across the voxels within each region, across stimuli within each category and across subjects (in a random-effects analysis). They observed that the activity within a region in the fusiform gyrus was stronger for face images compared to images depicting objects from other categories. Regions selective for other categories, including places [6] and bodies [7], have also been discovered with this approach.

**Figure 1:**
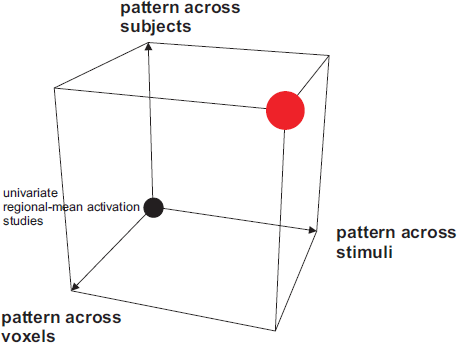
Honouring the unique response of each individual brain, in each cortical patch or column to each particular stimulus. Early neuroimaging studies have focused on analyses of regional-mean activation, averaging across voxels within the region of interest (e.g. a face region), across stimuli within a given category (e.g. individual faces) and across individual subjects (black circle). The field has begun to honour these distinctions, by analysing patterns of activity within each region, responses to single particular images and the variation across individuals. However, only recently have all these components of progress been combined (red circle) in a single study.

The approach of analysing three-way average activation has served the purposes of revealing the big picture, focusing analyses on what is consistent across individuals, simplifying the inference and increasing power by combining the evidence. However, averaging across individuals, cortical patches and stimuli also discards a lot of information in the brain-activity data that will ultimately be essential to understanding brain function.

An emerging literature is beginning to honour the idiosyncrasies that are at the heart of how the individual brain gives rise to the individual mind, endowing a complex world with a unique meaning and producing successful behaviour. We sketch a framework for multivariate analyses that link stimulus, brain representations and behaviour without averaging across stimuli, brain locations or individuals. In the context of this framework, we review previous studies in object vision that have made forays along these dimensions of progress. Finally, we describe what elements are still missing for cognitive neuroscience to engage the challenges of individuality.

## Representations: linking neurons to the computational goals of the brain

The concept of representation is central to the brain and cognitive sciences. Researchers often refer to a characteristic pattern of neuronal activity that reliably occurs when a particular stimulus is presented as a ’representation’ of the stimulus. When we refer to an activity pattern as a representation, we go beyond a statement of the statistical dependency between stimulus and response pattern established by data analysis. The term representation’ implies a functional interpretation, namely that the brain-activity pattern in question serves the function of representing the stimulus in the context of the organism’s overall brain information processing [8, 9]. This functional interpretation, although questionable in each particular case, has been extraordinarily helpful in building theories of brain function.

David Marr famously proposed the pursuit of brain science at three levels of description [10, 11]. The highest level is that of the computational goals of the system. The intermediate level is that of representations and algorithm. And the lowest level is that of neuronal implementation. This framework continues to guide theoretical and empirical brain science [12]. The representational interpretation, thus, provides the link between neuronal activity and the computational goals of the brain.

Cognitive psychology has tested cognitive theories with behavioural data, linking theory to experiment at a high level of description. At the opposite end of the spectrum, computational neuroscience has tested single neuron computational models with activity recorded from single neurons, linking theory and experiment at Marr’s lowest level. Both approaches are limited: behavioural data do not provide sufficient empirical constraints to explain brain function, and single-neuron computational models will never explain complex cognitive processes. Neuronal representations reside at an intermediate level that promises to link neurons to cognition. Neuronal representations are commonly associated with the activity of populations of neurons within a functional area. They exist at a spatial scale that lies between the level of single neurons, whose activity is classically recorded with electrodes, and the regional average activation of functional regions, which has been characterised by classical brain imaging methods. A major current challenge is to link theory and experiment at this crucial intermediate level of description, the level of neuronal population representations (Figure 2).

**Figure 2:**
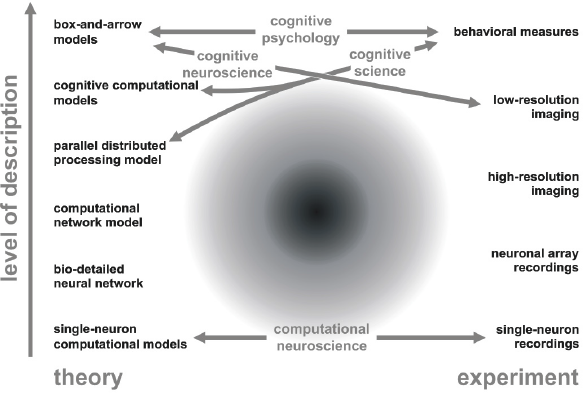
Levels of description in theory and experiment. The explanatory gap between single neurons (bottom) and cognition is bridged on the side of theory (left) by models at multiple levels of description. On the experimental side (right), the gap is bridged by neuronal array recordings and high-resolution imaging. However, the cognitive and brain sciences have yet to learn how to connect theory and experiment at the crucial intermediate level of description. The study of representational geometry offers one avenue of addressing this challenge.

In the domain of visual perception, the recognition of an object takes place along a hierarchy of visual areas [13], whose representational content ranges from local image features to representations of the parts of natural objects and their relationships, and on to semantic properties, as information moves forward along the ventral stream [14, 15, 16]. The cascade of representations arises through feedforward, lateral recurrent and feedback signalling between interconnected regions. Higher cognitive processes make use of the representations to produce successful behaviour and construct or update the organism’s knowledge [17]. Understanding the representation within each area would provide a major stepping stone towards understanding the brain as a whole.

## From univariate selectivity to pattern information

Brain science has experienced a paradigm shift from univariate analyses of selectivity to multivariate analyses of pattern information. This paradigm shift started with the theoretical concept of a neuronal “population code” and has more recently led to widespread multichannel measurement and multivariate pattern-information analyses (Figure 3) in both cell recording and functional imaging [18].

**Figure 3:**
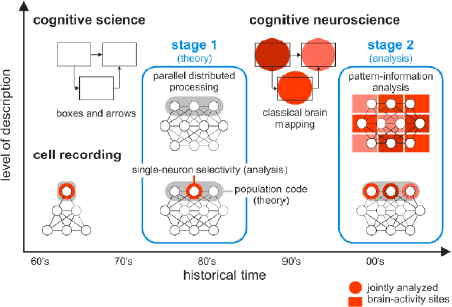
Population-coding theory and pattern-information analysis: a two-stage paradigm shift for systems neuroscience. The paradigm shift from univariate selectivity to pattern information in neuronal population codes had two historical stages. In the 1960s, cell recording studies mostly used single electrodes to measure single-neuron (or single-site) response amplitudes. The responses were analysed univariately to assess neuronal selectivity. In theory, similarly, the dominant idea was that information important to the animal should be explicitly represented in single-cell responses. Cognitive science proceeded quite separately from neurophysiology in this period, constructing box-and-arrow models based on behavioural data. Stage 1 of the paradigm shift concerned theory and occurred when population coding became the prevalent concept in neurophysiology. Cognitive science concurrently embraced the concept of parallel-distributed processing, which brought it closer to brain science. Although distributed population coding was the dominant theoretical concept, measurement and analysis of neuronal responses (red) remained mostly univariate. Then the advent of neuroimaging led to the inception of cognitive neuroscience. However, spatial resolution was initially low (in the cm range with positron emission tomography) and so the initial approach was to link brain regions to the boxes of the box-and-arrow models of cognitive psychology. Researchers assessed each region’s overall involvement in different tasks with univariate analyses. More recently, cell-array recordings and high-resolution functional imagining have enabled us to measure distributed representations in unprecedented detail. Stage 2 of the paradigm shift concerned analysis and occurred when the field began to measure large numbers of responses simultaneously within a region and to analyse them jointly with multivariate pattern-analysis techniques.

In neuroimaging, the paradigm shift towards pattern information analyses was pioneered by Haxby et al. [19], whose pattern-decoding approach revealed widely distributed category information in the ventral temporal cortex. This study was among the first to honour the distinct contribution to the representation of each little patch of cortex measured by an fMRI voxel. However, like previous work on category selectivity, response patterns were averaged across many distinct stimuli (which were presented in a category-block design), and although activity patterns were not averaged across individuals, individual idiosyncrasies were not investigated.

The approach described in Haxby et al. [19] was an example of the important concept of late combination of the evidence [20]. Combining the evidence across multiple measurements is essential when dealing with noisy data, as it improves the signal-to-noise ratio. The combination of the evidence can be achieved early on in the analysis procedure, by averaging activation levels across voxels, stimuli or individuals. However, averaging is not the only way to combine the evidence. Haxby et al. (2001) did not average across voxels within a region. Instead, they correlated category-related response patterns across voxels as part of their pattern-decoding approach. Pattern-information analyses, such as multivariate decoding, combine the evidence across voxels without averaging the activity levels. This exploits a major strength of fMRI, the large number of response channels (voxels), and can greatly enhance sensitivity [21, 4]. The case for late combination of the evidence has been articulated for evidence distributed across voxels [19, 4] across stimuli [20, 22] and across individuals [23, 19].

## From category averages to rich sets of particular stimuli

Following Haxby et al. [19], multivoxel decoding became popular in neuroimaging [24, 21, 25]. Most studies took a simple-decoding approach, asking, for example, whether regional activity patterns contain information about a particular stimulus dichotomy. While engaging the complexity of distributed representations, the literature largely ignored individual differences and seldom analysed the representation of individual stimuli. Kriegeskorte and Bandettini [4] investigated the pattern representations of particular stimuli. However, in order to obtain stable estimates of the response patterns, the study was limited to four particular object images, two faces and two houses.

A number of studies have explored more complex stimulus spaces, while averaging across voxels and focusing on commonalities across subjects [26, 27, 28]. Mur et al. [26] investigated whether the category preferences of regions in the visual ventral stream held for every exemplar of a set of 96 object images. They found that face and place regions exhibit almost perfectly categorical ranking of the single-image activations, but also graded responses within the preferred and non-preferred categories. This study also took a step towards honouring individually unique representations by investigating both subject-average and subject-unique activation profiles.

Three fMRI studies published in 2008 explored the pattern representations of richer sets of particular stimuli. Mitchell et al. [29] investigated the representation of noun concepts. They showed that a semantic model fitted on the basis of response patterns elicited by 58 word picture pairs could predict response patterns elicited by novel noun concepts (not used in training the model). Kay et al. [30] took a similar approach in the domain of vision, investigating the representation of particular images in early visual cortex. They showed that a Gabor-filter model fitted on the basis of response patterns elicited by 1750 training images could predict response patterns elicited by novel images.

Kriegeskorte and Bandettini [4] investigated the representation of 92 object images in the inferior temporal cortex of humans (hIT) and monkeys (IT). They found that the patterns associated with individual images formed clusters corresponding to natural categories and that these clusters (along with the within-category representational dissimilarities) matched closely between human and monkey. These three studies, reviewed in [31], all honoured the distinct contributions of individual voxels and the representations of particular stimuli. By exploiting the late combination of evidence, they managed to forgo averaging across stimuli, while exploring richer sets of stimuli than previous studies.

## From pattern information to representational geometry and tests of computational models

The three papers just mentioned [30, 22, 29] also took the analysis of representational patterns in a novel direction. While previous pattern analyses used generic statistical models to demonstrate the presence of information about the stimuli in a brain region, these three studies tested computational models of brain information processing, which predicted not merely the presence of information about particular stimulus dimensions, but the format in which the information was represented. In population receptive-field modelling [32, 30, 29], computational models are used to predict the responses of individual voxels. In representational similarity analysis [20, 22] computational models predict the dissimilarity relationships of the response patterns (Figure 4 and Figure 5). The fMRI activity pattern elicited by each particular stimulus is compared to the pattern elicited by each other stimulus. All pairwise comparisons are assembled in a representational dissimilarity matrix (RDM). RDMs are useful because they capture not only the information present, but also the format in which it is represented. Moreover, RDMs from brain regions can be directly compared to RDMs predicted by computational models. Population receptive-field modelling, by contrast, requires a separate data-set for fitting a linear model that predicts the response of each voxel from the computational model representation. RDMs also enable straightforward comparisons between brain regions, between individuals, between species and between modalities of brain-activity measurement [9].

**Figure 4:**
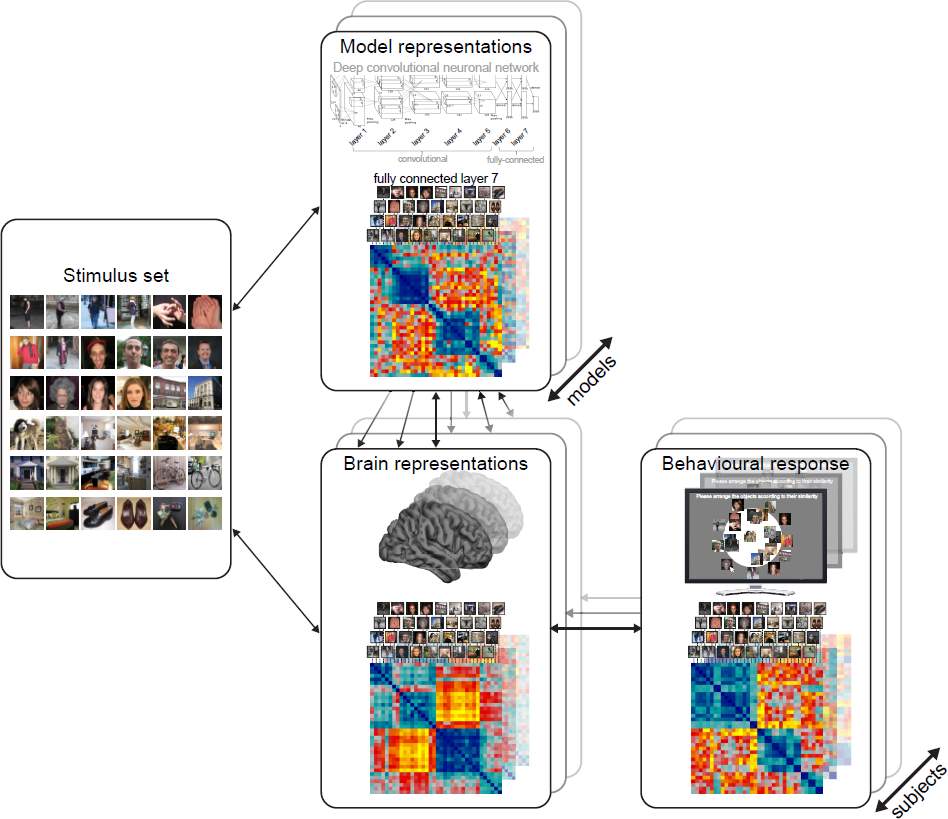
Combining multivariate descriptions of stimuli, brain representations, and behavioural responses. With recent advances in pattern information analyses, progress has been made in understanding object-vision processes in the brain, and to relate them to behaviour. Advances in computer vision enable us to model the similarity structure of stimulus properties with ecologically valid, neuropsychology inspired and biologically plausible computational modeed and biologically plausible computational models. One example of such computer vision model is the deep convolutional neuronal network (DCNN) model of vision, which made significant advances in object recognition and offer similar representational performance to human inferior temporal (hIT) cortex. For example, there is a great deal of correspondency in representational geometry between the fully connected layer 7 of the DCNN and the representational geometry in individual subjects hIT cortex. Future research will seek to establish whether understanding the computations achieved throughout the model’s architecture can help us understand the computational mechanisms that enable an individual’s brain to recognise objects. These brain representations can also be compared across subjects, and related to multidimensional accounts of subjective behaviour.

**Figure 5:**
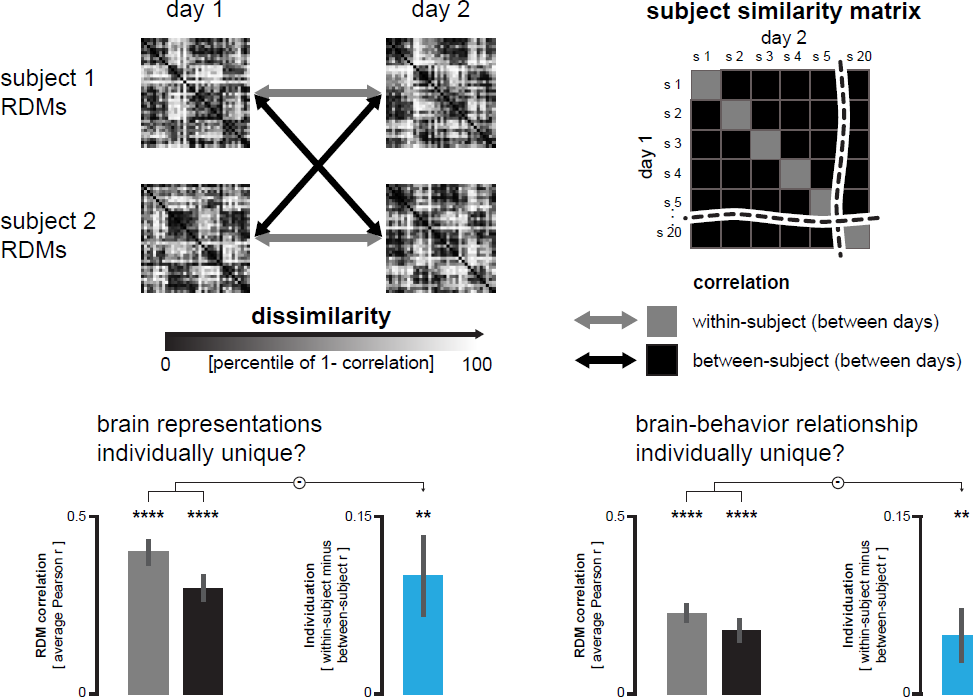
Characterising individually unique object representations. (a) Individual subjects are scanned on two separate days. This allows characterising their object representational geometry independently on each occasion. This allows characterising the stability of a representational geometry within and between subjects (red and black bars). This also enables us to compute an individuation index, consisting of the average within-subject RDM correlation across days, minus the average between-subject RDM correlation across days. If the within-subject RDM correlation is significantly larger than the between-subject RDM correlation, this indicates some degree of individual uniqueness in the representation. After defining whether components of the brain representations are individually unique, one can also investigate whether the brain representations are predictive of the idiosyncrasies in behaviour using the above mentioned framework. (b) Actual results from a recent study on individually unique brain representations. The left panel shows the stability of the representational geometry of the hIT activity patterns to visual objects (within-and between-subjects; red and black bar). The within-subject RDM correlation was significantly larger than the between-subject RDM correlation across days, reflecting the idiosyncrasies in the hIT representation (blue bar). The right panel shows a significant correlation between the brain representations (averaged across the two scanning days) and the similarity judgments obtained from the multiple arrangements task (within-and between subjects; red and black bar). The brain-behaviour correlation was significantly larger within than between subjects, indicating that an individual’s similarity judgments are better predicted by that individual’s mental representations than by another’s.

## From stable to task-flexible representations

The representational geometry of a brain region has recently been shown to be somewhat influenced by top-down mechanisms such as attention and behavioural goals [33, 34, 35, 36]. C¸ ukur et al. [33] used fMRI to study the impact that searching for an object category during a natural movie has on the semantic representation measured in the brain. The volunteers were asked to either “search for humans” or “search for vehicles” while their brain activities were recorded. Similarly to the procedure described in Huth et al. [37], the authors used WordNet to label object and action categories in natural movies. Using voxel-wise modelling and regularised regression, the authors showed how attending one category distorted the semantic structure of the neural representations of both attended and unattended categories, with category-attended expansion of the representational geometry at the cost of a compression of the distant category [33]. In another study, Harel et al. [34] used fMRI to investigate how the neural representations in regions of the ventral temporal cortex vary as a function of task. They compared the brain-activity patterns for a single set of objects under six different tasks (fixation, colour, tilt, content, movement and size). Their results demonstrated the presence of flexible task-dependent neural representations in the lateral occipital and in the lateral prefrontal cortex, indicating that object processing is highly influenced by the aim of the observer [34]. These two studies clearly demonstrated the task-adaptive flexibility of the representations of the ventral temporal cortex and of the prefrontal cortex. Both these studies were designed to characterise commonalities across individuals in the brain’s ability to flexibly adapt to task demands.

## From group characterisations to individually unique representations

Honouring individual voxels and stimuli, the cited studies have made important advances. However, most of this literature tacitly assumed that representations are consistent across individuals [38, 39, 24, 37, 30, 40, 22, 29, 41, 42, 43, 44, 45, 46]. Several studies have explicitly demonstrated representational commonalities between individual [19, 47] and even between species [22].

Even if all individuals had functionally identical brains and the spatial layout of functional units were similar across individuals, the precise anatomical location of functional units in a common brain space might still be variable. This would reflect limitations of our alignment methods (defining the common brain space) rather than true functional differences between individuals. Brains are commonly aligned using volume-based (Talairach or MNI) or surface-based (FreeSurfer, BrainVoyager, AFNI-SUMA) methods or individual functional localiser experiments.

Considering the variability of functional localisation across individuals (as it appears in a given common brain space) can improve the outcome of activation analyses. For example, Fedorenko and colleagues have stressed the importance of defining functional regions in a subject-specific manner, before averaging activation across subjects [48, 49]. Neuroimaging has a long history of individually defined functional regions of interest (e.g. Kanwisher et al., 1997). In pattern-information studies, similarly, activity patterns are usually not averaged across individuals [19, 21]. A recent study demonstrated that structure-function relationships can be consistent between individuals, even when the precise localisation in a given common brain space is variable. Saygin et al. [50] demonstrated that the precise location of face-selective responses in the fusiform gyrus of an individual subject can be predicted on the basis of anatomical connectivity measured with diffusion-weighted imaging. Face-selective responses in the fusiform gyrus were associated with a particular fingerprint of anatomical connectivity across the rest of the brain. The prediction model was cross-validated across subjects, demonstrating the consistency across individuals of the relationship between structure (anatomical connectivity) and function [50].

The most widely used paradigm of group analysis in neuroimaging assumes some common brain space and treats across-subject variation in activation levels as a random effect. In contrast to fixed-effects analysis, where the variability across subjects is not modelled at all, random-effects analysis models interindividual variation as noise [51, 52, 53]. It is important to note that neither of these approaches treats the interindividual variation as an effect of interest.

Despite the widespread focus on commonalities, there is also an expanding neuroimaging literature on individual differences (for a review, see [54]), with most studies addressing individual differences in regional-average activation. Individual differences in regional-average activation are commonly investigated in memory research, higher order cognitive functions, intelligence research and social neuroscience [55, 56, 57, 58].

As an example from the object-vision literature, Furl et al. [55] showed that regional-average face responses predict a person’s ability to recognise faces. Other studies have investigated how interindividual variability in connectivity [59] and brain anatomical measures relates to behavioural differences between individuals [60, 61, 62].

Most studies that investigated individual differences in brain function focused on regional-average activation and its relationship to behaviour. While pattern-information studies usually allow for the precise location of representational units to vary across individuals, few studies to date have addressed hypotheses about the individuality of brain representations. Can brain representational idiosyncrasies explain a person’s unique perception and behaviour?

Raizada et al. [63] investigated interindividual differences in brain representations and the degree to which they predict interindividual differences in perception. Using pattern-information analyses, they compared brain representations of the phonemes /ra/ and /la/ in English and Japanese speakers, showing that the behavioural ability to discriminate these vocalisations could be predicted from the discriminability of their representational patterns in auditory cortex [63, 64]. This relationship held across groups (English vs. Japanese speakers) and even across individuals within the two groups.

Another study investigated the role of the hippocampus in episodic memory using pattern-information analyses [65]. The authors showed that the activity patterns observed in the hippocampus carried information about the temporal positions of objects in learned sequences. Furthermore, individuals who performed better in object sequence retrieval had more robust hippocampal object-position binding as indicated by larger hippocampal voxel pattern similarity. The studies by Raizada et al. [63] and Hsieh et al. [65] used pattern representations measured in each subject to predict a unidimensional subject covariate (behavioural performance). A recent study by Charest et al. [23] investigated the relationship between high-dimensional perceptual judgements of the similarity among a set of objects and the multivariate brain representations of the objects. Earlier work has shown that object similarity judgements exhibit similar categorical divisions and similar within-category similarity structure as ventral temporal representational patterns [66]. Charest et al. [23] showed that even individual idiosyncrasies of the perceptual judgements could be predicted from idiosyncrasies of the brain representational geometries. The methodological approach and key result of this paper are shown in Figure 5.

## From univariate to multivariate descriptions of stimuli and behavioural responses

We have argued that brain representations are inherently multivariate and should thus be described and analysed multivariately. The same is true for the objects in the external world and our behavioural responses to them (Figure 6). The study by Charest et al. [23], cited above, demonstrated how we can investigate the individually unique relationships between multivariate brain representations and multivariate behavioural measures. The multivariate behavioural measure used in this study was similarity judgements. Judgements for all pairwise object comparisons were acquired using a novel multi-arrangement task [67, 68]. In this method, subjects communicate perceived object similarity by arranging object images on a computer screen by mouse drag-and-drop operations. In an adaptive measurement procedure, the subject is asked to arrange subsets of the objects in 2D according to their similarity. The higher dimensional perceptual similarity space is then inferred from the multiple arrangements by inverse MDS [68].

**Figure 6:**
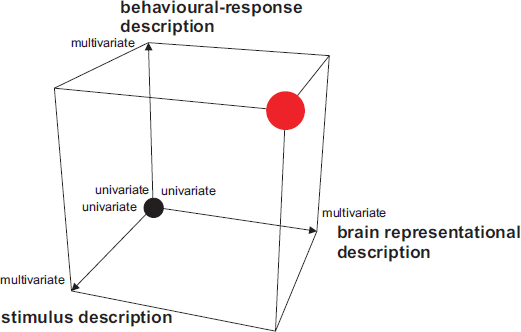
Multivariate description of stimulus, brain representation, and behaviour. Early studies have focussed on particular dimensions of stimulus, brain response, and behaviour. However, the sensory information, its representation in the brain, and behaviour are fundamentally multivariate phenomena. Recent studies have begun to engage this inherent complexity, linking stimulus to brain response, brain response to behaviour, or all three elements, while treating several of them in a multivariate framework.

Recent studies have increasingly employed multivariate descriptions of the stimuli. One example cited above is the study by Mitchell et al. [29], who used a 25-dimensional semantic space to describe their stimuli. A more recent example is the study by Huth et al. [37] who defined detailed semantic predictors describing a movie stimulus. The predictors described the presence of objects from different categories in the scene and utilised the is-a hierarchy of the WordNet model [69]. Using a regularised linear regression on the voxel activity patterns, they showed how the continuous semantic space provided by the WordNet labels predicted the activity of each voxel in the brain [37].

The semantic models of Mitchell et al. [29] and Huth et al. [37] require the stimuli to come with labels. The computational model linking stimuli to brain representations, thus, is not fully explicit in these studies. The cited study by Kay et al. [30] provides an example of a test of a simple image-computable model (Gabor model). Characterising the multivariate nature of the stimuli in the external world is now also possible for higher level representations, using more complex computational models. One influential model of object vision is the HMAX model [70, 71]. HMAX is a computational model inspired by the neuroscience literature, which aims at characterising the representations along the visual hierarchy. Efforts have been made to compare the representational geometry of the HMAX model to the representational geometry of activity patterns along the visual ventral stream [72, 20, 22, 73]. The classical implementation of the HMAX model so far fails to satisfactorily explain the categorical divisions of higher level object representations.

Recent advances with deep convolutional neural networks (DCNNs) have improved the performance of computers at visual object recognition [74, 75, 76]. These models are loosely biologically inspired in their hierarchical layered architecture and acquire their representations through supervised learning with large category-labelled image sets. Recent studies suggest that the performance achieved by the DCNNs in object classification approaches that of IT cortex in human and non-human primates [77]. The DCNN developed by A. et al. [74] outperformed other computer vision models at predicting the representational geometry of human and monkey IT [78] for isolated object images. These authors showed that the models strong categorical boundaries (acquired through strong supervision) contributed to its better performance at predicting the IT representational geometry. For the contextualised images from Charest et al. [23] as well, we observed that the A. et al. [74] model captures major categorical distinctions similarly to human IT (Figure 4, previously unpublished data). These recent studies have given us a better understanding of the computational mechanisms underlying object vision.

## Honouring the idiosyncrasies

Figure 7 explores the extent to which a number of exemplary studies conducted over the past two decades honoured the idiosyncrasies of voxels, stimuli and subjects. We see a clear development from averaging to more sophisticated data modelling that engages these idiosyncrasies. Future studies will push further along these dimensions, linking multivariate descriptions of the world via explicit computational models of brain information processing to multivariate measurements of brain activity in individual organisms and to multivariate measurements of behaviour. One framework for pursuing these directions is the analysis of representational dissimilarity matrices (Figure 4 and Figure 5; for a review, see [9]), which enable us to relate multivariate descriptions of stimuli, brain regions, model representations and behavioural responses without complex fitting procedures.

**Figure 7:**
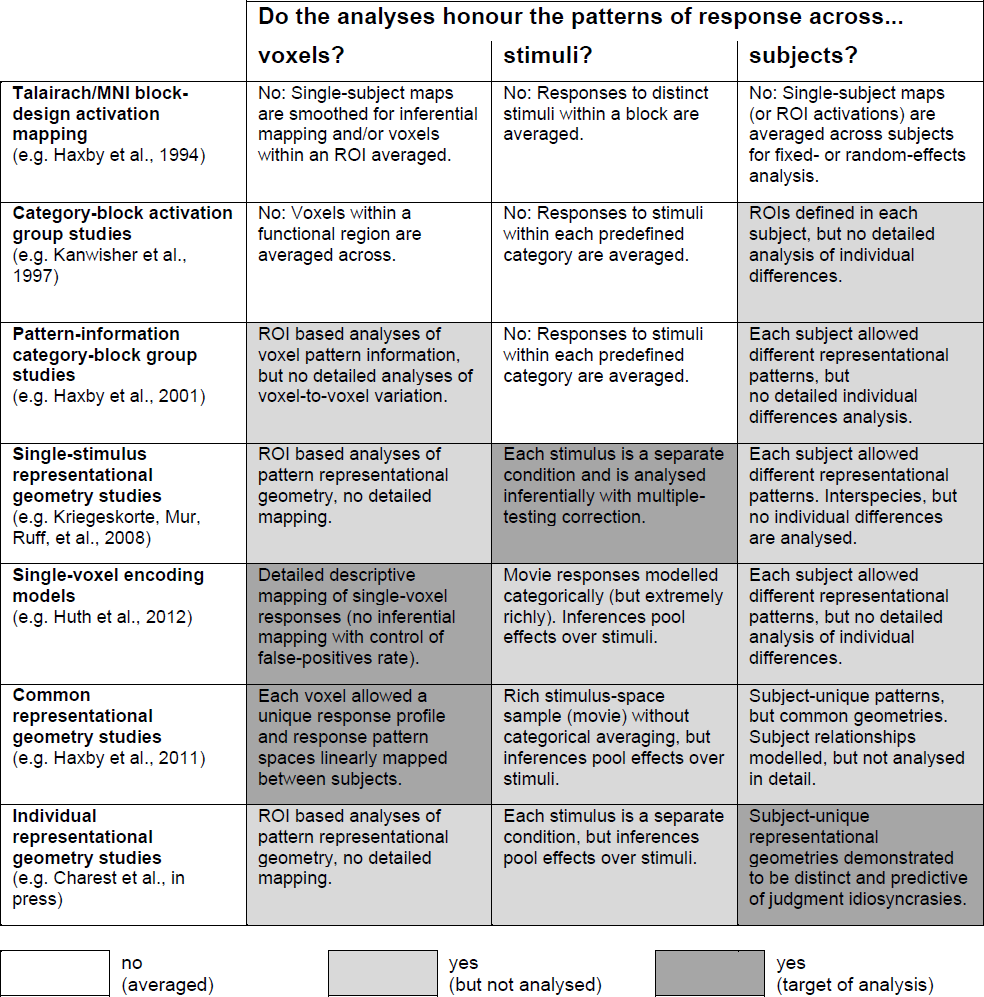
Honouring idiosyncrasies of voxels, stimuli, and subjects. Early neuroimaging studies (top row) tended to average across voxels, stimuli, and subjects. These studies gave us a coarse-scale view of brain function, revealing how the regional-average activation of large chunks of brain differed between tasks (each involving complex processing of particular stimuli, which were represented by block averages). The resulting brain maps were typically averaged across subjects. Recent studies have attempted to honour the idiosyncrasies of voxels, stimuli, and subjects. Here we compare particular exemplary studies (cited in the left column) with regard to the way they treated variation across these dimensions. We distinguish two levels of honouring differences between voxels, stimuli, and subjects: (1) The differences are modelled and not treated as noise, but they are not the target of the analyses (light gray). The motivation for this approach is usually the greater power such analyses can confer for testing hypotheses of interest. For example, pattern-information studies that do not average activation across voxels or subjects tend to have greater power for detecting information encoded in subject-unique patterns of activity within a region. (2) The differences are not only modelled, but form the target of the analyses. An additional more technical dimension that we do not consider here is the distinction between fixed- and random-effects analyses (across subjects, stimuli, or voxels). An analysis that treats subject as a random-effects factor is useful for generalising from the sample of subjects to the population. However, the variation across subjects is part of the noise model (thus not honoured) and the inference targets the population mean.

Beyond basic science, the characterisation of individually unique brain function is likely to contribute to our understanding of disorders and of the continuous variation across patients. If functional brain imaging is to become useful in the diagnosis and monitoring of patients, we will need to develop a rich repertoire of methods for characterising subtle functional differences between individual brains and minds. Disorders are increasingly recognised to fall on a complex manifold where every patient has a unique place. For example, individuals with an autism spectrum condition (ASC) tend to exhibit atypical perceptual processing [79]. However, every case is different. Grandin and Panek [80] describe three challenges that neuroimaging researchers of autism face: the absence of apparent brain structural abnormalities [81], the heterogeneity of causes of ASC and the heterogeneity of behavioural symptoms. In the future, it might become possible to characterise the biological underpinnings of the unique behavioural dysfunction of a given individual patient. Multivariate characterisations of an individual’s brain representations might one day help tailor therapeutic interventions in the context of personalised medicine.

## Acknowledgement

We would like to thank Philip Pell for helpful comments on the manuscript.

